# Role of a single MCP in evolutionary adaptation of *Shewanella putrefaciens* for swimming in planktonic and structured environments

**DOI:** 10.1101/2024.10.15.618407

**Authors:** Daniel B Edelmann, Anna M Jakob, Laurence G Wilson, Rémy Colin, David Brandt, Frederik Eck, Jörn Kalinowski, Kai M Thormann

## Abstract

Bacteria can adapt to their environments by changing phenotypic traits by mutations. However, improving one trait often results in deterioration of another one, a trade-off which limits the degree of adaption. The gammaproteobacterium *Shewanella putrefaciens* CN-32 has an elaborate motility machinery comprising two distinct flagellar systems and an extensive chemotaxis array with 36 methyl-accepting chemotaxis sensor proteins (MCPs). In this study we performed experimental selection on S. *putrefaciens* for increased spreading through a porous environment. We readily obtained a mutant that showed a pronounced increase in covered distance. This phenotype was almost completely caused by a deletion of 24 bp from the chromosome, which leads to a moderately enhanced production of a single MCP. Accordingly, chemotaxis assays under planktonic conditions and cell tracking in soft agar showed that the mutation improved navigation through nutritional gradients. The study demonstrates how differences in the abundance of a single MCP can lead to an efficient upgrade of directed flagella-mediated motility in specific environments at a low expense of cellular resources.

**Importance:** Experimental evolution experiments have been used to determine the trade-offs occurring in specific environments. Several studies that have used the spreading behavior of bacteria in structured environments identified regulatory mutants that increase the swimming speed of the cells. While this results in a higher chemotaxis drift, the growth fitness decreases as the higher swimming speed requires substantial cellular resources. Here we show that rapid chemotaxis adaptation can also be achieved through modification of the chemotaxis signal input at a low metabolic cost for the cell.

## Introduction

Many bacteria are motile by means of flagella, which are long helical proteinaceous filaments that extent from the cell surface and are rotated by a cell envelope-embedded motor. Flagella allow swimming through liquid environments and swarming across appropriate surfaces (Berg and Anderson, 1973; Kearns, 2010; Wadhwa and Berg, 2021). To allow navigation towards or away from certain environmental conditions and to get access to nutrients, bacterial flagellar motility is controlled by one or more chemotaxis systems (Matilla *et al*., 2023). Environmental signals are perceived by different methyl-accepting chemotaxis proteins (MCPs), which assemble to large arrays within the cytoplasmic membrane. The signals sensed by the MCP affect the activity of the chemotaxis histidine kinase CheA, which phosphorylates the response regulator CheY. CheY∼P directly interacts with the flagellar motor(s) to induce switches in the rotational direction or, in some cases, pausing of rotation, which change the direction of movement by various mechanisms. By adjusting the run lengths between reorientation events according to the cues sensed by the MCPs, the cells modulate their otherwise random walk and can thus actively move up or down a gradient of these cues (Sourjik and Wingreen, 2012; Bi and Sourjik, 2018; Grognot and Taute, 2021; Thormann *et al*., 2022).

The formation and operation of one or more flagella is metabolically costly and results in a significant growth disadvantage of flagellated cells under uniform culturing conditions, where motility does not provide an advantage (Berg, 2003; Martínez-García *et al*., 2014; Ziegler and Takors, 2020; Schavemaker and Lynch, 2022). Therefore, regulation of motility gene expression has evolved in a way to optimize the fitness trade-off between the metabolic burden and the benefit of chemotaxis and motility in dependence of the corresponding environmental conditions (Taylor and Stocker, 2012; Yi and Dean, 2016; Fraebel *et al*., 2017; Ni *et al*., 2017, 2020; Colin *et al*., 2021). Accordingly, an elaborate gene network regulates the expression and formation of bacterial flagella and chemotaxis systems as a function of environmental cues (Amsler *et al*., 1993; Chevance and Hughes, 2008; Guttenplan *et al*., 2013; Prüß, 2017). A well-studied system for such a regulation is the carbon catabolite repression of *Escherichia coli*, which shows that flagella-mediated motility and chemotaxis become more important in environments were carbon sources are scarce (Adler and Templeton, 1967; Amsler *et al*., 1993; Liu *et al*., 2005; You *et al*., 2013; Hui *et al*., 2015; Ni *et al*., 2020). Thus, *E. coli* follows a growth strategy when carbon sources are plentiful but switches to a search strategy when the carbon source is scarce (Ni *et al*., 2020; Colin *et al*., 2021).

Experimental evolution experiments on bacteria with respect to enhanced chemotaxis have shown that the balance between growth and motility can be readily shifted by mutations. In the cases that have been studied in more detail, enhanced chemotaxis could be mainly attributed to an elevated expression of flagellar genes, which resulted in a higher swimming speed and, by this, an increased chemotactic drift (Yi and Dean, 2016; Ni *et al*., 2017; Colin *et al*., 2021).

*Shewanella putrefaciens* CN-32 is a facultatively anaerobic gammaproteobacterium. Compared to *E. coli*, which harbors a peritrichous flagella and a single chemotaxis system with five MCPs, the flagella and chemotaxis machinery of this species is considerably more intricate. *S. putrefaciens* possesses two distinct flagellar systems, a polar and a lateral system, which are encoded in two separate gene clusters (Bubendorfer *et al*., 2012). The monopolar flagellum is the primary system that is formed under most conditions and mediates main propulsion, screw thread motility, and chemotactic responses (Bubendorfer *et al*., 2012; Kühn *et al*., 2017). The chemotaxis system is located to the flagellated cell pole (Rossmann *et al*., 2015) and has an extensive signal perception range with 36 putative methyl-accepting chemotaxis sensor proteins (MCPs). The secondary flagellar cluster leads to the formation of one to five lateral flagella whose motors do not respond to the chemotactic system. They solely turn counterclockwise and do not form a bundle like the paradigmatic *E. coli* flagellar system. The additional flagella provide additional thrust while moving through structured environments and lower the turning angles during directional switches of the cells during chemotaxis. By this, they positively affect spreading in structured and liquid environments (Bubendorfer *et al*., 2012, 2014; Kühn *et al*., 2022). Their formation is induced in a subpopulation of cells when complex nutrients are available and likely under conditions of high load on the main polar flagella. However, the regulatory mechanisms underlying the formation of lateral flagella system is still mostly obscure (Schwan *et al*., 2022).

Previous studies have demonstrated that for *S. putrefaciens* efficient motility and spreading in structured environments such as soft agar, depends on numerous factors. These include the nutrient content of the medium, the cells’ chemotactic ability, the geometry of the polar filament and the presence of lateral flagella. In addition, the c-di-GMP-dependent production of adhesion factors, such as the MSHA type 4 pili, has an influence on spreading (Bubendorfer *et al*., 2012, 2014; Kühn *et al*., 2017, 2018, 2022; Pecina *et al*., 2021; Rick *et al*., 2022). Considering the more intricate flagellar set-up of *S. putrefaciens* as compared to that of *E. coli*, we asked whether an experimental evolution towards better spreading would yield regulatory mutants affecting the polar and/or lateral flagellar system. To answer this question, we evolved mutants with enhanced spreading capability in soft agar. Instead of differences in global regulation of flagellation, we identified a mutation that leads to a slightly increased production of a single MCP and the resulting increased chemotaxis in the selection environment as the almost exclusive reason for the observed pronounced increase in spreading. Thus, rapid chemotaxis adaptation can be achieved through a modification of chemotaxis signal perception at a low metabolic cost for the cell.

## Results

### S. putrefaciens soft-agar spreading evolution

A number of different factors can affect *S. putrefaciens* spreading in structured porous environments. To get more insights into the importance and regulation of the different factors, e.g. the regulation of the secondary lateral flagellar system, we adopted an experimental evolution approach. We aimed at isolating spontaneous mutants of *S. putrefaciens* that are better able to spread through soft agar, a common assay to determine motility and chemotaxis ability of bacteria. To monitor easily via fluorescence microscopy the flagellation status, a possible target of our selection pressure, we used for the spreading evolution approach an *S. putrefaciens* CN-32 strain that allows coupling of maleimide-ligated fluorescent dyes to both the polar and lateral filament. For dye coupling, the flagellin-encoding genes were mutated for the flagellins to harbor serine-to-cysteine substitutions at surface-exposed positions on the flagellar filament (Kühn *et al*., 2022). Within this study, this strain will be referred to as wild type.

The procedure of the experimental evolution is depicted in **Figure 1A**. Cells from an exponentially growing culture were spotted on a LB soft-agar plate and were allowed to spread for 24 h. Cells were then isolated from the fringes of the visible spreading zones and re-inoculated on fresh LB soft agar. After 14 repetitions, the isolated cells exhibited a highly increased spreading diameter (**Fig. 1B, C**). The up-motile phenotype was stably retained after taking the strain into stock and culturing on solid or liquid media. Thus, we considered the mutation genetically fixed. The up-motile strain was referred to as *S. putrefaciens* G14.

**Figure 1:**
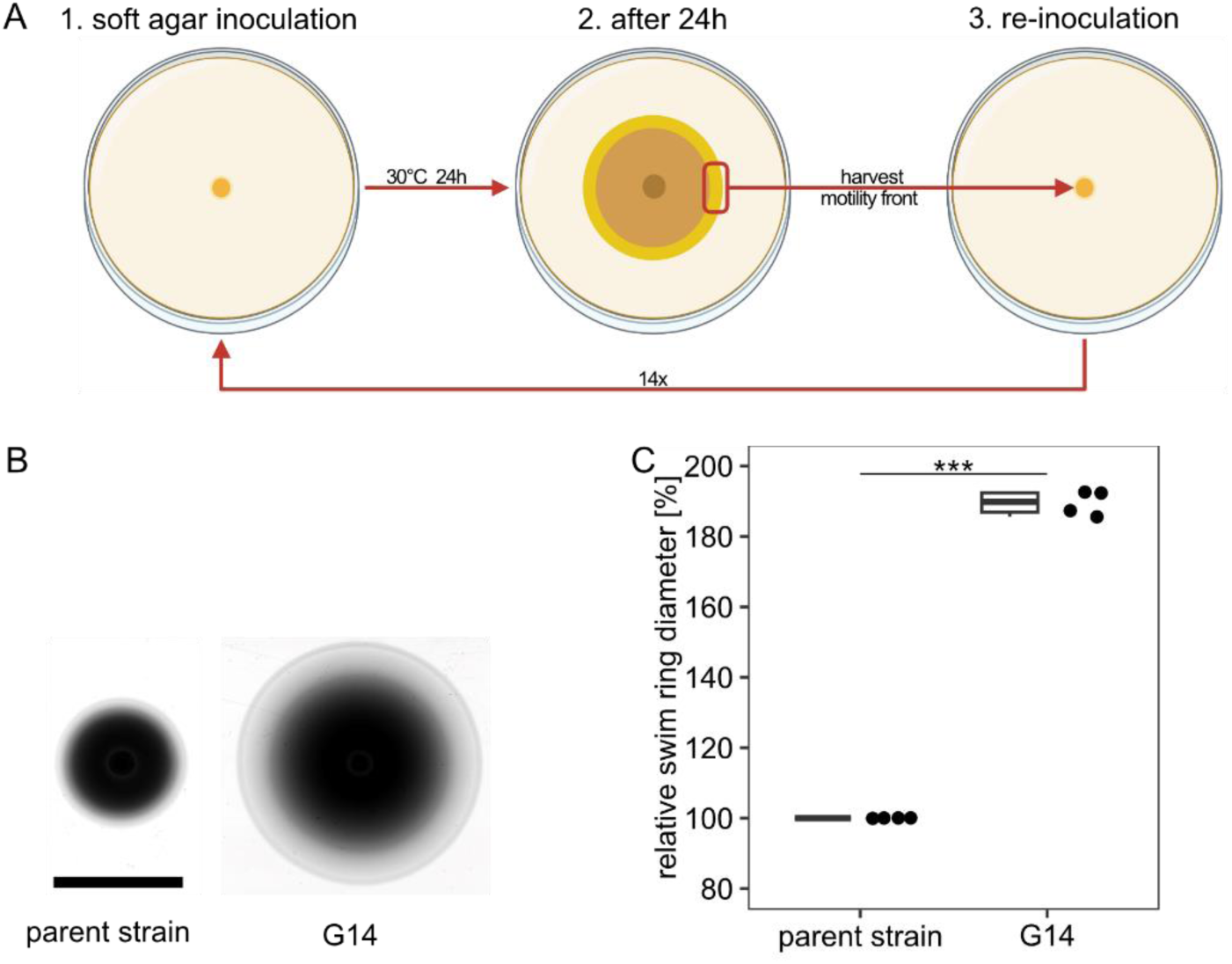
Isolation of an *S. putrefaciens* CN-32 up-motile mutant by experimental evolution. **A)** Procedure of isolation. An *S. putrefaciens* strain was allowed to spread on LB soft agar for 24 h. Cell material was isolated from the outer fringes of the spreading zone and re-inoculated on soft agar. The procedure was repeated 14 times. **B)** Comparison of the parent and the evolved strain (G14) on soft agar. **C)** Quantification of the spreading radius of 4 independent experiments. The asterisks indicate the significance according to a pairwise.t.test (p < 0.001).

Time-lapse scans show that the evolved strain G14 instantly outperforms the wild type in spreading, and constantly outpaces the wild type as spreading zones grow (see **Supplementary Movie 1**). Therefore, we assumed that the advantage in soft-agar motility is present at inoculation and does not develop at a later stage of spreading.

### Up-motile mutant cells are not affected in flagellation and morphology

Previous studies showed that production of lateral flagella and the length of the main polar filament affect spreading of *S. putrefaciens* in soft agar (Bubendorfer *et al*., 2014; Kühn *et al*., 2017, 2018). As the next step, we therefore determined the flagellation state of wild-type and G14-mutant cells. To this end, cells were isolated from planktonic cultures and soft-agar plates. The flagellar filaments were fluorescently labeled and then used for microscopic observation. The cell morphology did not differ between mutant and wild-type cells, and we observed no significant difference with respect to the main filament length (**Figure 2; Supplementary Figure 1**). The number of flagellated cells remained similar as did the average flagellar number per cell, and there was no apparent difference in flagellation pattern of lateral flagella. We therefore conclude that mutations affecting the general regulation of flagellation are unlikely.

**Figure 2:**
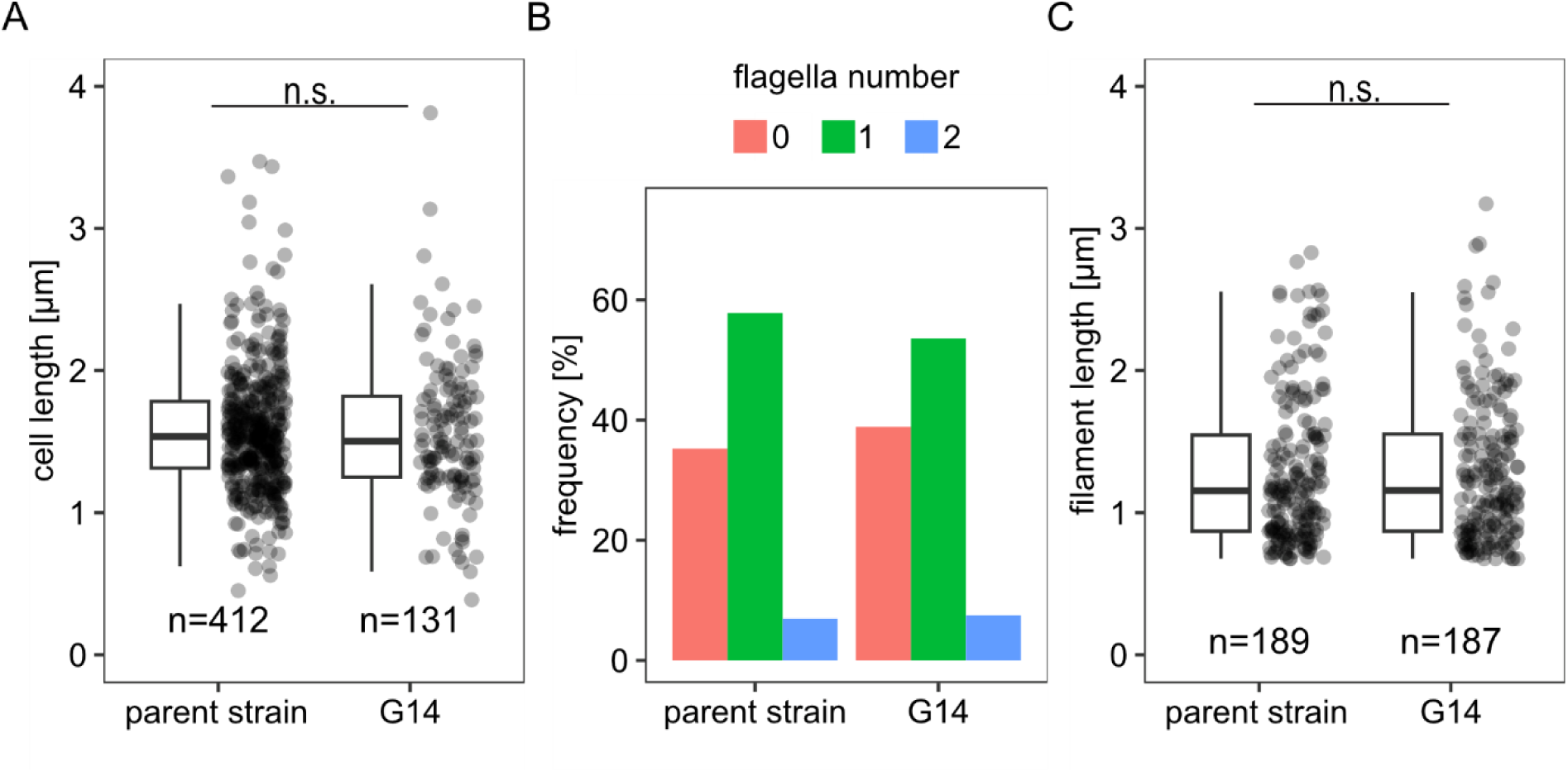
Cell morphology and flagellation of the G14 mutant. Quantification of **A)** length of cells, **B)** percentage of flagellated cells **C)** length of filaments, each in liquid media. n.s., not significant according to a pairwise.t.test, the number of cells is indicated by n.

### Identification of a single MCP as the main cause of increased spreading

To identify the nature of the mutation underlying the increase in spreading, three different clones of the evolved up-motile strain G14 were sequenced. Apart from the cysteine substitutions in the flagellins, the only mutation occurring in all sequenced mutants was a deletion of 24 bp upstream of the orphan gene Sputcn32_0387 (**Figure 3A**). The predicted encoded protein is 638 amino acids in length, has a molecular mass of 69 kDa and represents one of the 36 MCPs encoded by *S. putrefaciens* CN-32. The N-terminal region (up to amino-acid position 287) of the MCP (from now on referred to as MCP_0387) is predicted to localize to the periplasm and possesses a Cache_3-Cache_2 domain, which likely serves in signal perception. The identity of the signal is currently unknown. Homology comparisons indicate that homologous MCPs are present in numerous other species of *Shewanella*. As the deletion upstream of MCP_0387 was also the only mutation that was apparently directly related to flagella-mediated motility, we focused on the role of this mutation in the up-motile phenotype.

**Figure 3:**
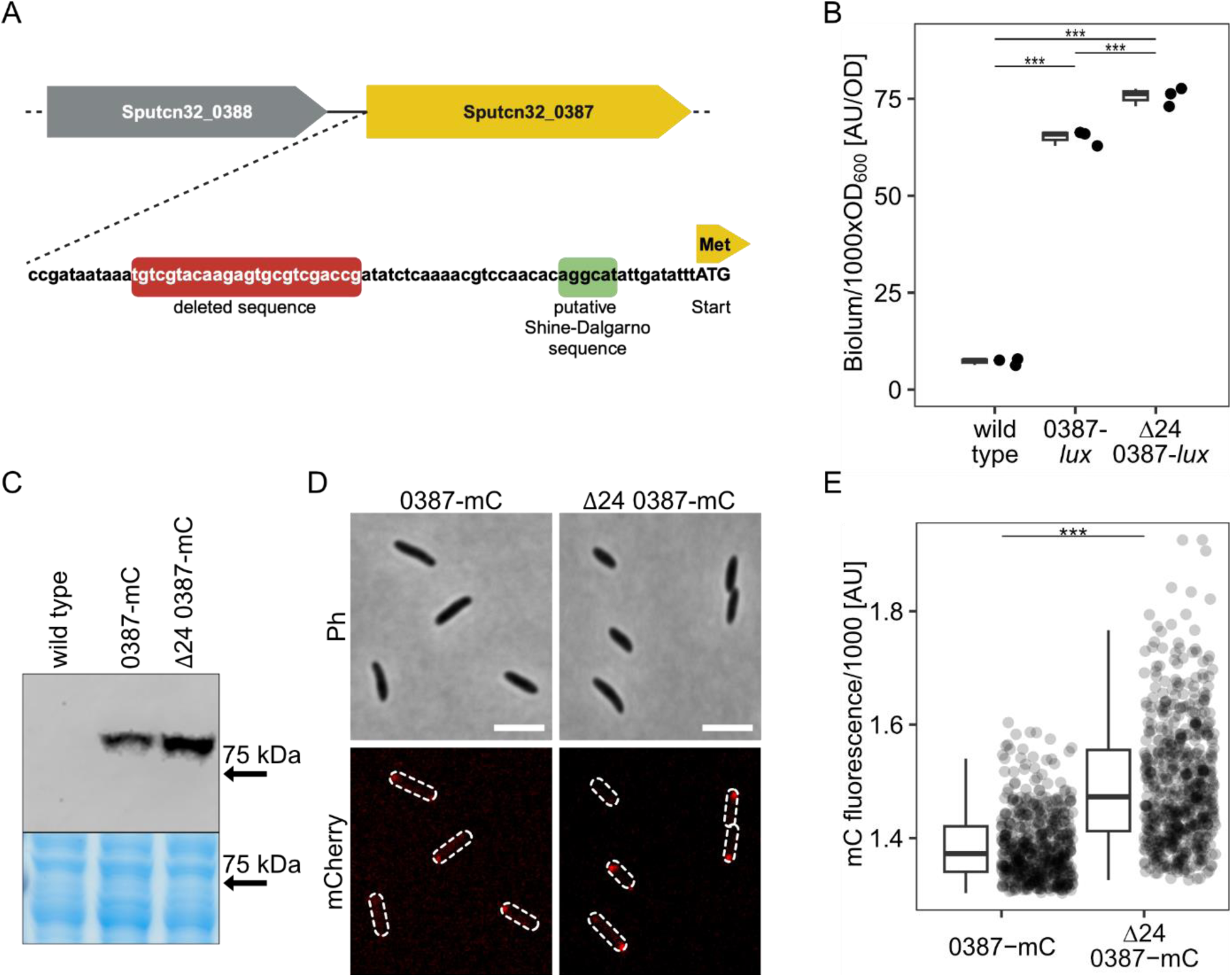
Mapping of the dominant mutation to MCP_0378. **A)** Cartoon of gene region. A 24-bp region was found to be deleted upstream of SO_0384, encoding an MCP (MCP_ 384). **B)** Quantification of SO_384 expression by integration of *luxCDABC* directly downstream of the structural gene. Light units emitted by the indictated strains were determined in exponentially growing cultures. **C)** Comparison of produced MCP_0387-mCherry in the wild-type and the Δ24 mutant backgrounds by western blotting. The panel below shows the same region of the Coomassie-stained gel prior to blotting as loading control. For the full uncut gel and western see Supplementary Figure 3. **D)** Localization of MCP_0387-mCherry by fluorescence microscopy. Shown are micrographs of cells expressing the hybrid fusion gene from the chromosome. The upper two panels show the phase contrast image, the lower two the corresponding mCherry fluorescence image where the position of the cells is outlined. The scale bar equals 5 µm. For an image including the untagged wild type see Supplementary Figure 4. **E)** Quantification of the polar mCherry fluorescence intensity. **B/D)** The asterisks display the significance using a pairwise.t.test (p < 0.01).

The deleted region located 37 bp upstream of the MCP-encoding gene’s start codon. To first determine whether MCP_0387 is involved in enhanced spreading through soft agar, we introduced an in-frame deletion of the corresponding gene in the evolved strain *S. putrefaciens* G14. Soft-agar assays showed that the up-motile phenotype was completely lost in this mutant (**Supplementary Figure 2**). This result indicated that MCP_0387 is directly involved in the observed spreading phenotype and that the mutation does not abolish the transcription of the MCP_0387-encoding gene.

For further analyses, we introduced a line of mutations into the *S. putrefaciens* wild type to exclude any potential effects of the other mutations we identified through sequencing. Fist, we constructed a double mutation including the Δ24 deletion mutation along with a deletion of the downstream gene encoding MCP_0387 (Δ24Δ0387). In this strain, Sputcn32_0387 was re-integrated into its native position on the chromosome in *S. putrefaciens* Δ24Δ0387 (*S. putrefaciens* Δ24). Soft-agar spreading assays using the three strains showed that the deletion of the 24 bases upstream of the MCP-encoding gene enhanced spreading of the mutant cells (**Supplementary Figure 2**). An additional deletion of MCP_0387 abolished the increase in spreading completely, and the strain displayed an even lower spreading than wild-type cells (86.9 %). Re-integration of the gene restored the increased spreading phenotype. Based on the results, we proposed that MCP_0387 was mainly responsible for the gain-of-function with respect to increased motility in soft agar.

### The identified deletion mutation increases the production of MCP_0387

The 24 bp gene deletion was located upstream of Sputcn32_0387 and deletion of the gene completely abolishes any advantage in spreading and even decreases spreading ability. Therefore, we assumed that, due to the deletion, the transcription activity is affected in a way that leads to a higher abundance of the MCP. To determine the promoter activity, we created a transcriptional fusion to a *lux*-based reporter by placing a *luxCDABE* gene cassette from *Photorhabdus luminescens* into the chromosome directly downstream of Sputcn32_0387. By this, the expression of Sputcn32_0387 could be directly quantified *in vivo* via luminescence emitted by the cells. Compared to the wild-type background, the luminescence of the Δ24 mutant was significantly but only moderately higher (about 116 %; **Figure 3B**). As the next step, we aimed at enabling visualization and quantification of MCP_0387. To this end, we created a Sputcn32_0387-mCherry hybrid gene, which was integrated into the appropriate strains to replace native Sputcn32_0387. This resulted in the production of stable MCP_0387 with a C-terminal fusion to the fluorophore mCherry. Fluorescence microscopy and western blotting showed that MCP_0387-mCherry is produced and polarly localized in all wild-type and Δ24 mutant cells, but the abundance of the MCP is moderately increased in the latter strain by a factor of about 1.5 (**Figure 3C-D; Supplementary Figures 3 and 4**). From this, we concluded that the deletion leads to a moderate increase in the production and abundance of the MCP, which in turn enhances spreading in soft agar.

### MCP_0387 affects chemotaxis of S. putrefaciens

The identification of an MCP as the major contributor for enhanced spreading in soft agar suggested that the observed phenotype is mediated by differences in signal perception during chemotaxis. To first determine if the increased spreading is not due to enhanced swimming speed rather than chemotaxis signalling, we performed 3D-tracking of wild-type, Δ24 and Δ24Δ387 cells under planktonic conditions. We found that the cells of the three strains displayed a similar distribution of swimming speeds and that the maximum speed (about 62 µm · s^-1^) is not different between the strains (**Figure 4A; Supplementary Figure 5A**). The distribution of run durations and tumble frequencies was also highly similar under these conditions (**Supplementary Figure 5B and C**).

**Figure 4:**
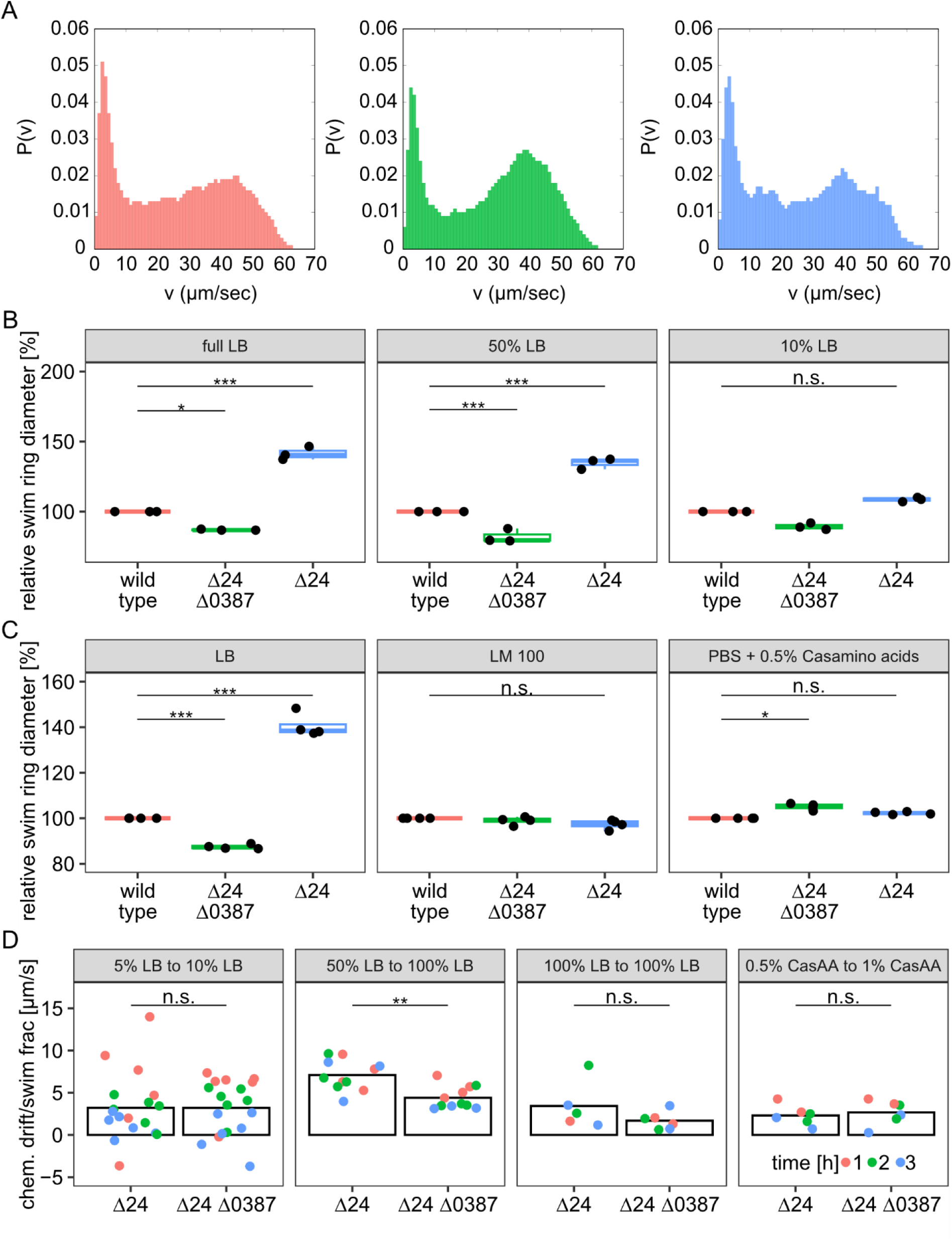
Effects of MCP_0387 levels on chemotaxis. **A)** Quantification of the swimming speed under planktonic conditions by 3D tracking. Shown is the speed distribution of the wild type (left panel; red; n = 281), the Δ24Δ0387 mutant (middle panel; green; n = 935) and the Δ24 mutant (right panel; blue; n = 265). **B)** Quantification of wild-type and mutant spreading at different LB conditions as indicated. Shown are the results from at least three biological replicates. **C)** Quantification of wt and mutant spreading under different media conditions (LB, LM and PBS buffer plus casamino acids) as indicated. Shown are the results from at least three biological replicates. **D)** Quantification of the chemotactic drift in controlled gradients as indicated. Shown are the results of at least three biological replicates after 1 (red), 2 (green) and 3 (blue) hours of incubation. The asterisks display the signifcance using a pairwise.t.test (*, p < 0.05; ***, p > 0.01; n.s., not significant).

Having ruled out swimming speed as the predominant factor for the observed increase in spreading, we reasoned that altering the chemotaxis response by different levels of MCP_387 may be dependent on the environmental nutrient conditions. Therefore, we next determined whether the up-motile phenotype is retained in media other than full-strength LB (**Figure 4B and C**). We observed that in soft-agar spreading assays with decreasing concentrations of LB the spreading advantage of the mutant was retained at 50% LB. In contrast, the spreading advantage became non-significant in 10% LB, in LM medium (a HEPES-buffered medium containing small amounts of yeast extract and peptone and lactate as main carbon source), or in a buffer with 0.5% casamino acids.

To disentangle the effects of growth, diffusion properties and chemotaxis, which combine to give rise to a given spreading ability on soft agar, we also probed the drift of the mutants up controlled gradients of the different media, using microfabricated chambers that connect two reservoirs with different concentrations of chemoeffectors via a small rectangular channel (Colin *et al*., 2014). For this, we used the Δ24 and the Δ24Δ0387 mutants, between which we expected the largest differences according to the soft-agar spreading assays. The cells start from the low concentration reservoir and drift towards the other reservoir by a combination of diffusive flux and chemotactic bias. The Δ24 and Δ24Δ0387 strains show similar drifts in absence of initial gradients in LB medium (**Figure 4D**), indicating that their diffusion is similar in a liquid medium. In contrast, the Δ24 strain shows a stronger drift in response to a large gradient of LB than the Δ24Δ0387 strain (**Figure 4D**). Conversely, both strains have similar drifts in a gradient of casaminoacids. Hence, the receptor 0387 is responsible for a specific chemotactic response to LB gradients that contributes to the enhanced spreading of the Δ24 strain on LB soft agar.

### MCP_387 levels affect motility in soft agar

To more accurately determine the movement patterns of the MCP_0387 mutant cells as compared to those of wild-type cells, we performed a holographic analysis of motility within the soft agar matrix. To this end, a soft-agar spreading assay was set-up and holographic imaging was performed at the migration front of the spreading halo at a region where the cell density allowed to obtain tracks of single cells. The wild-parent, the Δ24Δ0387 and the Δ24 were used for imaging. We observed that within the soft agar, the cells exhibited periods of frequent directional changes with no or only small overall progress, which were interrupted by longer runs (**Figure 5A**). Such a ‘hop-and-trap’ motility has been observed previously for flagellated bacterial cells entrapped in a polysaccharide matrix (Wolfe and Berg, 1989; Bhattacharjee and Datta, 2019b, 2019a; Grognot *et al*., 2021, 2023). We then performed quantification of cells’ trajectories to determine differences in soft-agar spreading between the three strains. The ‘confinement ratio’ for the wild type and mutants, defined as the ratio of a cell’s net displacement to the total path length over the course of its track, were remarkably consistent between strains. The same was true for the direction correlation (**Supplementary Figure 6**).

**Figure 5:**
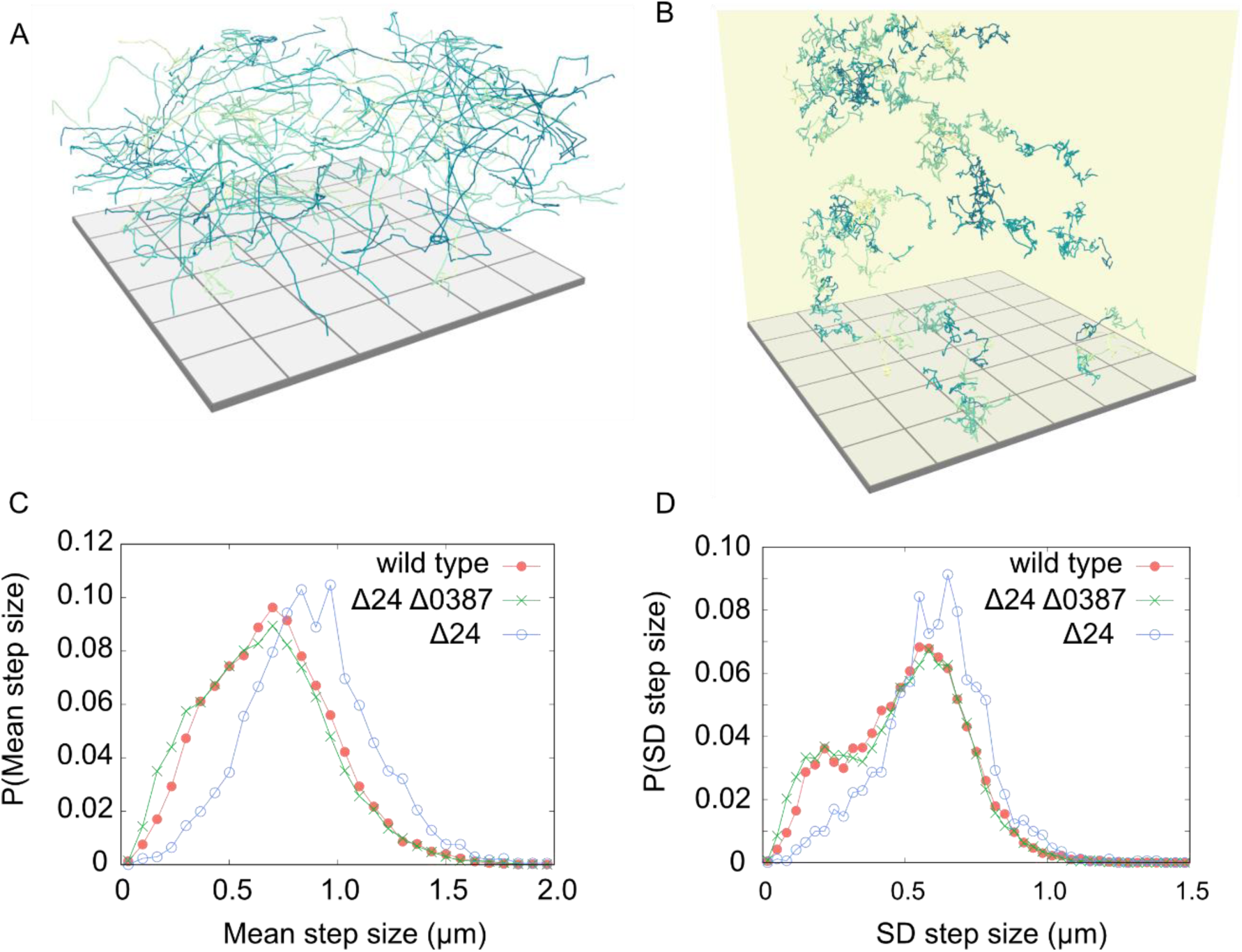
MCP_0387 affects spreading in soft agar. 3D-track projection of wild-type cells under planktonic conditions **(A)** and in soft agar **(B)**. The side of each square at the bottom is 100 µm. **C)** Quantification of the step size of the indicated cells in soft agar (wild type, red; Δ24Δ0387, green; Δ24, blue). **D)** Quantification of the standard deviation (SD) of the step size (wild type, red; Δ24Δ0387, green; Δ24, blue). The number of tracks quantified was 9,931 for the wild type, 11,393 for the Δ24Δ0387 mutant and 1,709 for the Δ24 mutant. Further parameters can be found in **Supplementary Figure 6**.

While confinement ratio and direction correlation were consistent between the strains, *S. putrefaciens* Δ24 mutant cells exhibited two important differences compared to the other two strains. First, the Δ24 mutant cells had, on average, longer runs (‘hops’) between the trap phases, as indicated by a shift of the mean step size towards higher values (**Figure 5**). In addition, the standard deviation (SD) of the Δ24 mutant step sizes was similarly shifted towards higher values. Together with the increase in step size, the results indicate that the cells tend to spend less time in the ‘trapped’ mode.

### MCP_387-enhanced spreading does not significantly affect growth

Previous studies showed that experimental evolution of soft agar spreading or swarming, which was based on increased expression of flagella genes and formation of longer or additional flagella, resulted in a significant decrease in growth under the same conditions. Therefore, we determined whether this would be similarly true for the increased spreading caused by MCP_387. Corresponding growth experiments showed no significant difference between the wild-type und mutant strains in nutrient rich (LB) medium and in nutrient-limited (4M) medium (**Figure 6**).

**Figure 6:**
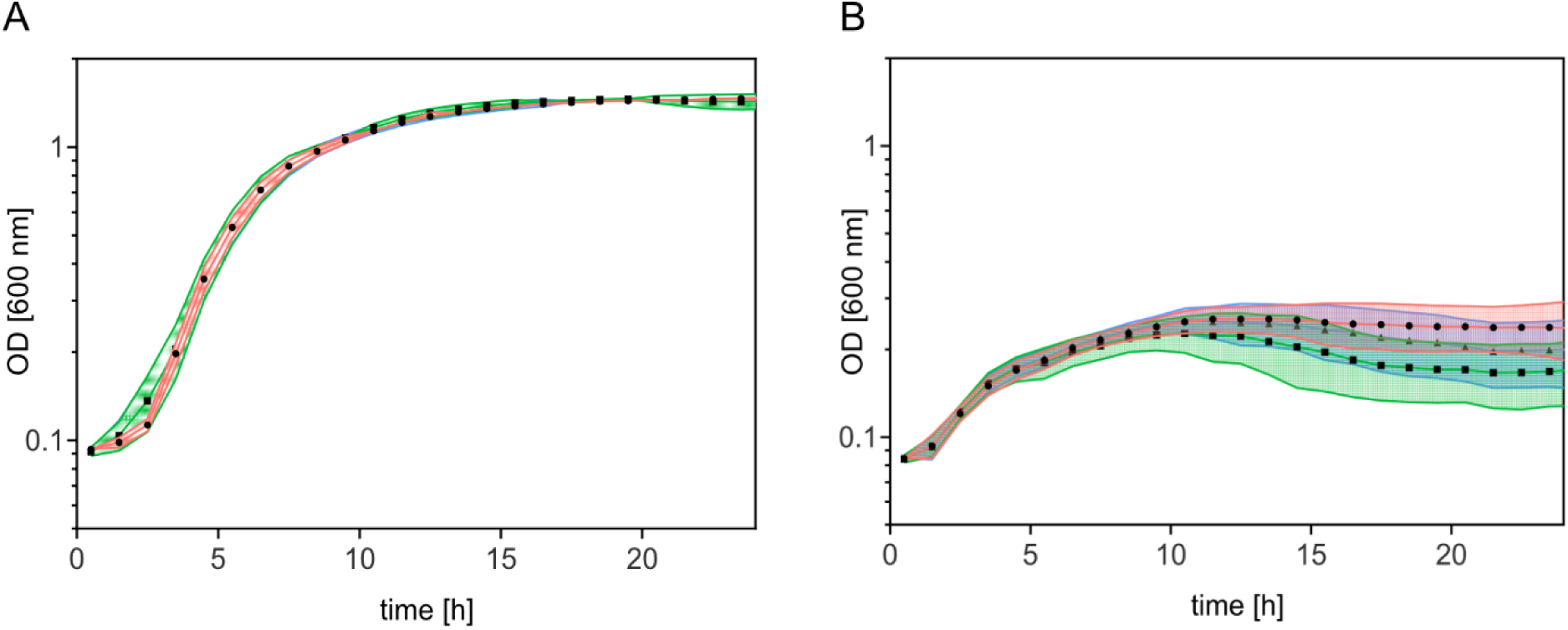
Effect of MCP_0378 levels on growth. Growth of the indicated strains in **A)** LB and **B)** 4M medium. Shown are the results of three biological replicates. The wild type is displayed in red, the Δ24 mutant in blue and the Δ24Δ0387 mutant in green.

## Discussion

Experimental evolution experiments on bacteria have emerged as a useful tool to study various aspects of adaptation and evolution (Fry, 2003; Perfeito *et al*., 2007; Kawecki *et al*., 2012; Taute *et al*., 2014; Ferenci, 2016; McDonald, 2019). Here we performed experimental evolution in soft agar using *S. putrefaciens* cells with a dual flagellar system to explore the potential of the underlying regulatory system for mutations beneficial to spreading in complex environments*. S. putrefaciens* shows complex regulatory interplay between the main polar and secondary lateral flagellar systems under these conditions. However, instead of a general flagellar regulation, a frequently observed evolution path in *E. coli*, we could attribute the better spreading of the up-motile mutant we obtained to overproduction of a single chemotaxis sensor protein, MCP_0387. Accordingly, our results demonstrated that the observed phenotype was due to enhanced chemotactic capability of the mutant strain under appropriate nutrient conditions.

Generally, flagella-mediated motility requires bacteria to balance the fitness trade-off between the benefits of chemotaxis and costs of allocating cellular resources to motility, and, accordingly, flagella synthesis is subject to intricate regulation at several levels. The trade-off between motility and growth was commonly observed when cells were experimentally selected for enhanced spreading and chemotaxis (Yi and Dean, 2016; Fraebel *et al*., 2017; Ni *et al*., 2017). The mutations causing the up-motile phenotype were usually mapped to factors directly or indirectly involved in flagella regulation. A common mutation that underlies a better spreading phenotype in *E. coli* strains is the integration of insertion sequence (IS) elements into the regulatory region of the *flhDC* operon, resulting in upregulation of flagella synthesis (Barker *et al*., 2004; Wang and Wood, 2011; Lee and Park, 2013; Zhang *et al*., 2017). Other studies on experimental evolution on *E. coli* spreading identified further regulatory key elements, such as the FlgM/FliA checkpoint, that govern flagella formation or length. These mutations affect the swimming speed and, by this, alter the chemotactic drift (Ni *et al*., 2017). Similarly, experimental evolution of a swarmer mutant of monopolarly flagellated *Pseudomonas aeruginosa* identified a mutant in the regulator of the flagellar number, FleN. The isolated FleN mutants form more polar flagella that are enabling better swarming (i.e. flagella-mediated movement across surfaces), but are negatively affected in growth (van Ditmarsch *et al*., 2013). In our case, the mutation did not occur in a regulatory factor but in the chemotactic sensory system, resulting in a highly environment-dependent upgrade of spreading. Despite its narrow medium specificity, this specialist evolution path does not require the expense of a large protein amount, as would be the case if more or longer flagella were synthesized, which limits the trade-off with respect to growth. As we performed an isolation of just one line of mutants, it is not clear if mutation of the chemotaxis system would be the predominant outcome of experimental evolution in *S. putrefaciens* under these conditions. It is possible that the more complex regulatory network of this species with its two flagellar systems does not as readily evolve as that of *E. coli*. Alternatively, a fine tuning of environment perception via receptor level changes might be easier for a species that has many more receptors than *E. coli*. More parallel lines of evolved *S. putrefaciens* strains under different environmental are required to address this question in future experiments.

MCP_0387 is only one out of 36 potential MCPs, which indicate a large chemotactic sensory repertoire of *S. putrefaciens*, and the level of its overproduction is rather small (a factor of 1.2 – 1.5). So how can such a small increase in MCP abundance have such a pronounced consequence? Previous studies showed that MCPs form trimers of dimers, which assemble into large well-structured arrays harboring the different MCPs (Yang and Briegel, 2020). The tight and organized clustering enables and facilitates cooperativity and signal processing, so that diverse signals can be processed and enhanced (Bray *et al*., 1998; Sourjik, 2004; Tu, 2013; Parkinson *et al*., 2015; Frank *et al*., 2016). We therefore assume that the increased abundance of one type of MCP, here MCP_0387, shifts the sensing capability and, correspondingly, the chemotactic response towards a yet unknown signal present in our selection media. Of note, a recent study on *E. coli* migration through soft agar demonstrated that cells isolated from the migration front tend to possess higher levels of the MCP Tsr, but only when the corresponding ligand serine was present (Vo *et al*., 2024). In contrast to the case described here for *S. putrefaciens* MCP_0387, the elevated Tsr levels occurred as the result of general heterogeneity in Tsr production in an *E. coli* population. Due to their ability to better chase the self-created traveling serine concentration gradient, these cells are then moving toward the front of the spreading zone (Fu *et al*., 2022; Vo *et al*., 2024). It is conceivable that such a heterogeneity in MCP levels similarly occurs in *S. putrefaciens*, albeit with 36 instead of just 5 MCPs, and that, by permanently increasing average levels, the mutation enriches the population in cells with high expression of MCP_0387 that may play a similar pioneering role very efficiently in our conditions.

Generally, the chemotactic potential of *Shewanella* sp. has only started to be explored and, so far, the signal the MCP responds to remains unknown. Notably, close homologs to MCP_0387 are widespread among *Shewanella* species, suggesting that the signal can be sensed and responded to by a number of these ubiquitously occurring bacteria. A potential orthologue of MCP_0387 in *S. oneidensis* MR-1, SO_4454, (92% identity/95 % positives) had been implicated as a potential candidate for energy taxis in this species, however, the effect was rather minor (Baraquet *et al*., 2009). The periplasmic domain of MCP_0387 harbors a Cache domain (Cache_3-Cache_2), which comprises the largest superfamily of extracellular sensors (Upadhyay *et al*., 2016). Accordingly, they have been implicated in binding a large range of potential ligands that may occur in the complex media containing tryptone and yeast extract, where the up-motile phenotype was visible. An analysis of the sensing MCP’s sensor domain did not give any direct hints with respect to potential ligands, and, so far, none of the potential ligands we tested (e.g., amino acids, small peptides) has given any clue (data not shown). The nature of signal that *S. putrefaciens* and likely other *Shewanella* species respond to via MCP_0387 and the corresponding potential orthologues is subject of current studies.

The chemotactic swimming pattern of bacteria at the single cell level has been mainly studied in detail during free swimming in planktonic environments. However, the movement pattern of single cells in structured environments differs substantially from that of their free-swimming counterparts (Adler, 1966; Armstrong *et al*., 1967; Wolfe and Berg, 1989; Croze *et al*., 2011; Amchin *et al*., 2022). Under planktonic conditions, the movement phases (‘runs’) are interrupted by short reorientation events. In contrast, when moving through soft agar, cells tend to be stalled for longer periods of time between runs. It is assumed that the cells are trapped in a pore within the polysaccharide matrix and require some reorientation events until an opening within the pore allows another run (Wolfe and Berg, 1989; Bhattacharjee and Datta, 2019b, 2019a; Grognot *et al*., 2023). The 3D-tracking in this study showed that *S. putrefaciens* cells perform a similar ‘hopping and trapping’ movement (Bhattacharjee and Datta, 2019a) in soft agar (**see Fig. 5A**). We found that the *S. putrefaciens* up-motile mutants are biased towards longer runs, as would be expected for an enhanced chemotaxis response. In addition, the mutant cells tend to spend a shorter time in the trapped mode. For *E. coli*, the likelihood of leaving the trap is thought to depend on successful reorientation of the cell, the ‘tumble’, which is affected by the geometry of the pore (Bhattacharjee and Datta, 2019a). In contrast, *S. putrefaciens* and other polarly flagellated species navigate by a run-reverse-flick pattern, and flagellar wrapping and lateral flagella activity may assist in leaving the trap (Kühn *et al*., 2017, 2022, 2022; Thormann *et al*., 2022; Grognot *et al*., 2023). In addition, diffusion and, therefore, local nutrient concentrations may be different under these conditions. Further studies are required to determine flagellar behavior and identify the exact mechanism underlying the shorter periods of the *S. putrefaciens* MCP_0387 mutant in the trapped state in soft agar.

## Materials & Methods

### Growth conditions and media

*E. coli* and *S. putrefaciens* CN-32 strains used in this study are listed in **Supplementary Table 1**. Unless specified otherwise, cells were cultured in LB medium at 30°C (*S. putrefaciens*) and 37 °C (*E. coli*). Kanamycin and 2,6-diaminoheptanedioic acid were used as selection markers at final concentrations of 50 mg ml^-1^ and 300 µM, respectively.

Growth experiments of *S. putrefaciens* strains were carried out in 1 ml LB or 4M (Gescher *et al*., 2008) medium at 30 °C in a 24-well microtiter plate using a BioTek Epoch 2 Microplate Spectrophotometer (Agilent). Each experiment was carried out in at least three biological replicates with four technical replicates each.

### Strain constructions

All *S. putrefaciens* modifications were constructed as outlined previously (Bubendorfer *et al*., 2012). In brief, double homologous recombination was applied using 500-bp flanking regions upstream and downstream of the target region in order to generate open reading frame deletions or integrate open reading frames with point mutations or genetic markers. Recombination was achieved using the suicide plasmid pNPTS-R6K after conjugation from E. coli WM3064 (Lassak *et al*., 2010) as listed in **Supplementary Table 2**. Vectors were constructed using standard Gibson assembly protocols (Gibson *et al*., 2009) using primer listed in **Supplementary Table 3**.

### Soft-agar spreading assays

Soft agar was prepared using 0.25% (w/v) select agar (Invitrogen) in LB medium, lactate medium (LM100) (Paulick et al, 2009) or PBS supplemented with 0.5% (w/v) casamino acids (Carl Roth). The soft-agar plates were inoculated with strains pre-grown to exponential growth phase and incubated at 30°C overnight. Strains to be directly compared were always inoculated on the same plate.

Time-lapse experiments of soft-agar spreading assays were performed using Epson Perfection V39 scanners. Image series were recorded using the scanlag software (Levin-Reisman *et al*., 2010). Each petri dish was inoculated with a single colony and incubated at 30°C for 18 hours while all assays were scanned in 15-minutes time intervals. Each image series was analyzed using Fiji ImageJ (Version 1.54i). To this end, the image series was converted to an image stack where the 0-minute image was used for background subtraction. Images were converted to 8-bit and a median filter (5 pixels) was applied. Colony outlines were segmented by applying the “threshold triangle” method and quantified by the “analyze particle” functions.

### Spreading evolution assay

For spreading evolution experiments, strain S8493 was used to perform repeated soft-agar spreading assays in 0.25% select agar in LB medium under standard conditions. Cells were harvested from the motility front by careful pipetting and were used to re-inoculate the consecutive spreading assay. After 14 cycles, cells harvested from the motility front were used to inoculate fresh LB medium to prepare cryo stocks in 10% DMSO to be stored at -80°C.

### Flagellar filament labeling and microscopy

In order to visualize flagellar filaments, surface exposed threonine resides of the flagellin monomers were exchanged to cysteine as outlined previously (Kühn *et al*., 2017). For microscopy, cells were harvested from soft-agar spreading assays by excising the agar fragment bearing the motility front and incubating it in PBS for 5 min. The cell suspension war separated from agar fragments, centrifuged (1200 × *g*, 5 min, room temperature) and resuspended in 50 µL PBS. Filament staining was performed by applying 2 µL Alexa Fluor 488 C5 maleimide (Thermo Fisher Scientific) for 18 min in the dark. Cells were washed with 1 ml PBS and sedimented by centrifugation for three times. Stained cells were applied to agarose patches (1% (v/v) select agar in PBS) for microscopy using a custom microscope setup (Visitron Systems) based on a Leica DMI 6000 B inverse microscope (Leica) equipped with a pco.edge sCMOS camera (PCO), a SPECTRA light engine (lumencor), and an HCPL APO ×63/1.4–0.6 objective (Leica) using a custom filter set (T495lpxr, ET525/50m; Chroma Technology). For phase contrast and fluorescence microscopy 50ms exposure times were used.

Cell body measurements were performed based on phase contrast images using the BacStalk software (version 1.8) (Hartmann *et al*., 2020). Filament measurements were performed using fluorescent microscopy images using Fiji ImageJ (Version 1.54i). Filaments were segmented on images using grey value 2000 as threshold for background subtraction and the “convert to mask” functions. Object lengths were measured using the “skeletonize (2D/3D)” and “analyze skeleton (2D/3D) functions.

### Strain sequencing

Chromosomal DNA was extracted using the E.Z.N.A Bacterial DNA kit (omega BIO-TEK) according to the manual instructions appropriate for Gram-negative bacteria. From extracted genomic DNA TruSeq PCR-free sequencing libraries (Illumina, San Diego, USA) were generated according to the manufacturers’ instructions and sequenced on an Illumina MiSeq machine (2x300bp) with V3 chemistry. Separate libraries were generated for the parental strain as well as two descending strains after 7 and 14 generations of selection. Reads were imported into Geneious Prime 2022.0.1 (https://www.geneious.com) and analyzed using the pre-defined “Map reads then find variations/SNPs” workflow. Detected variants against the *S. putrefaciens* CN-32 reference (GenBank: CP000681.1) were then exported for further filtering and analysis.

### Chemotaxis experiments in liquid medium

Chemotactic drift measurements were performed similarly to previous works with *E. coli* (Colin *et al*., 2019) and *V. cholerae* (Irazoki *et al*., 2023). In short, PDMS microchambers were constructed by molding Sylgard 184 (Dow Corning) mixed in 10:1 base-to-crosslinker ratio on a SU8-based Silicon wafer template, curing overnight at 65°C, peeling off and cutting the hardened PDMS to shape, and binding it to a glass slide via oxygen plasma treatment. The chambers were filled with DI sterile water 20 minutes after production to preserve the hydrophilicity of their surfaces until same-day use. The microchamber consists of two large reservoirs connected by a small channel (LxWxH = 2mm x 1mm x 70 µm). The first reservoir receives a first solution of indicated composition, which contains bacteria at 10^8^ – 10^9^ cells/mL concentration, while the second reservoir is filled with a second solution, which is initially cell-free. After filling the device and sealing it, a gradient of concentrations forms in the channel, which reaches a steady state profile in 1h in absence of consumption. We measure the drift of the cells in this gradient 1, 2 and 3h after sealing in the middle of the channel at mid-height under a phase contrast microscope (Nikon *Ti*) with a 10x (NA 0.3) objective and a CMOS camera (Mikrotron Eosens CXP, 1px = 1.4µm) that records a movie at 200 frames/s for 100 s.

The movie is analyzed using dynamic differential microscopy (DDM) (Wilson *et al*., 2011), to measure swimming speed v0 and fraction of swimming cells φ, and phase differential microscopy (ϕDM) (Colin *et al*., 2014), to measure population averaged drift v_d_ as described previously. The algorithms are implemented as publicly available ImageJ Plugins (Colin *et al*., 2019). In DDM, the differential intermediate scattering function is computed from the spatial Fourier components of the images. It is then fitted with a model that accounts for a mixed population of diffusing non-motile cells and of motile cells, which are modeled as 3D swimmers with swimming speed taken from a Schultz distribution (Wilson *et al*., 2011). The average swimming speed v0 and the fraction of swimmers φ are then determined as the average value of the fitted parameter on the range of wave numbers where the fit converges (typically, q = [0.5, 2.0] px^-1^). In ϕDM, the mean drift R(t) of the population of cells is extracted from the shift in the phase of the Fourier components as described previously (Colin *et al*., 2014). A linear fit of the drift over the whole movie yields the population averaged drift velocity v_d_. Since the drift of non-swimmers is negligible, the (chemotactic) drift of the motile cells is estimated as v_ch_ = v_d_/φ.

### Holographic microscopy data acquisition

Digital holographic microscopy was configured as described previously (Kühn *et al*., 2018; Thornton *et al*., 2020; Findlay *et al*., 2021). In brief, a single-mode optical fiber was held in the condenser mount of an inverted microscope, and the light directed down on to the sample stage. Samples were imaged using a 20× magnification objective lens and a Mikrotron MC-1362 camera, resulting in a final magnification of 0.70 µm per pixel. Videos were acquired at a resolution of 1024×1024 pixels, and frame rates of 10 Hz or 20 Hz for cells embedded in agar, and 100 Hz for the planktonic analysis.

### i) Planktonic samples

Tracking experiments in liquid medium were performed with cells during exponential growth phase diluted to an optical density of 0.1 at 600nm with fresh LB medium. The samples were contained in chambers constructed from glass slides and UV-curing glue to provide a volume measuring approximately 5×20×0.3 mm^3^. Samples were loaded by capillary action from one end and sealed with petroleum jelly to prevent evaporation. Three movies were acquired from each strain. The movies were detrended by subtracting a fitted 4^th^ order polynomial function to the intensity series of each pixel; this removes static background artifacts, as well as any slow drifts in the sample, but maintains the images of the motile cells. Each frame of these detrended movies yielded a three-dimensional reconstruction of the optical field within the sample. The field was reconstructed using Rayleigh-Sommerfeld back-propagation (Lee and Grier, 2007), creating stacks of numerically reconstructed images from each raw data frame. The optical field in the sample was reconstructed in around 100 axial planes uniformly separated by 3 μm throughout the sample volume, and image stacks were segmented based on axial intensity gradients (Wilson and Zhang, 2012; Farthing *et al*., 2017). Each video frame was processed independently to allow parallelization, with coordinates in subsequent frames linked together across time to make cell tracks. Motile and non-motile cells were distinguished using their mean-squared displacements (MSDs). Objects were identified as non-motile and discarded if their average MSD per unit time, or average squared displacement at one second were too low. Tracks were regularized using piecewise cubic splines to obtain better estimates of instantaneous swimming speed. Cell reorientations (‘tumbles’) were identified as peaks in angular velocity, and the time between consecutive reorientation events taken to be the duration of one run. Tracks must contain at least two reorientation events to contribute to the run duration statistics.

### ii) Agar samples

For 3D holography tracking experiments of cells in agar, all strains were cultured under standard conditions as outlined for soft-agar plates. The samples were imaged on an inverted microscope in the same manner as the planktonic ones, with the plate lid removed while the video sequences were obtained. Videos Detrending of the video sequences to remove background artifacts was performed as in the planktonic case. The optical field was reconstructed using 250 axial slices of 3 μm, giving a total sample volume of 720×720×750 um^3^ within the agar, but the post-processing was otherwise the same as in the planktonic case.

### Statistical analysis

Statistical tests were performed using the pairwise.t.test function of R statistical language (version 4.2.2). For calculations, two sided, non-pooled standard deviations, unpaired were set as parameters. P-value adjustments were calculated according to the Benjamini-Hochberg correction method.

## Acknowledgements

This work was supported by a grant from the Deutsche Forschungsgemeinschaft DFG (TH831/8-1) to KMT. RC acknowledges support from DFG grant CO1813/2-1.

