## Supplementary Tables 1-3 and Supplementary Figures 1-6; Caption Supplementary Movie 1 for "Role of a single MCP in evolutionary adaptation of *Shewanella putrefaciens* for swimming in planktonic and structured environments"

Affiliations:
[1]Institut für Mikrobiologie und Molekularbiologie, Justus-Liebig-Universität Gießen, Gießen, Germany
[2]Department of Physics, University of York, Heslington, York, YO10 5DD, UK.
[3]Max Planck Institute for Terrestrial Microbiology, and Center for Synthetic Microbiology (SYNMIKRO), Marburg, Germany
[4]Center for Biotechnology, Bielefeld University, Bielefeld, Germany

### Supplementary Material

Supplementary Tables 1 – 3
Supplementary Figures 1 – 6
Caption to Supplementary Movie 1

**Supplementary Table 1: Strains used in this study**

| Strain | Genotype | Source/Reference |
|---|---|---|
| *Escherichia coli* | | |
| DH5α-λpir | φ80d*lacZ* ΔM15 Δ(*lacZYA-argF*)U169 *recA1 hsdR*17 *deoR thi*-l *supE*44 *gyrA*96 *relA*1/*λpir* | Miller and Mekalanos, 1988 |
| WM3064 | *thrB*1004 *pro thi rpsL hsdS lacZ*ΔM15 RP4-1360 Δ(*araBAD*) 567*ΔdapA* 1341::[erm pir(wt)] | W. Metcalf, University of Illinois at Urbana–Champaign, IL |
| *Shewanella putrefaciens* CN-32 | | |
| S271 | CN-32 wild type | |
| | CN-32 Δ$flaAB_1$ Δ$flaAB_2$ | Bubendorfer *et al.*, 2012 |
| | CN32 Δ$flaA_1$_T174C $B_1$_T166C_S174C Δ$flaAB_2$ | this study |
| | CN32 $flaA_1$_T174C $B_1$_T166C_S174C $flaA_2$_T159C $B_2$_T159C | this study |
| | 14G clone 5 (evolved strain after 14 repeats) | this study |
| | 14G clone 9 (evolved strain after 14 repeats) | this study |
| | 14G clone 13 (evolved strain after 14 repeats) | this study |
| | 14G clone 5 Δ24Δ0387 | this study |
| | 14G clone 9 Δ24Δ0387 | this study |
| | 14G clone 13 Δ24Δ0387 | this study |
| | Δ24ΔCDS_0387 | this study |
| | Δ24 | this study |
| | CDS_0387-LuxCDABE | this study |
| | Δ24 CDS_0387-LuxCDABE | this study |
| | CDS_0387-mCherry | this study |
| | Δ24 CDS_0387-mCherry | this study |

**Supplementary Table 2: Plasmids used in this study**

| Name | Properties | Reference |
|---|---|---|
| pNPTS138-R6KT | *mobRP4*⁺ *ori*-R6K *sacB*; suicide plasmid for in-frame deletions; $Km^r$ | Lassak *et al.*, 2010 |
| pNPTS138-R6KT_Δ$flaA_1$_T174C $B_1$_T166C_S174C | insert for flagellin gene modification in pNTPS138-R6K | this study |
| pNPTS138-R6KT_$flaA_2$_T159C $B_2$_T159C | insert for flagellin gene modification in pNTPS138-R6K | this study |
| pNPTS138-R6KT_Δ24ΔCDS_0387 | insert for introduction of Δ24ΔCDS_0387 deletions in pNTPS138-R6K | this study |
| pNPTS138-R6KT_Δ24_CDS_0387knock-in | insert for re-insertion of Δ24ΔCDS_0387 in Δ24 in pNTPS138-R6K | this study |
| pNPTS138-R6KT_CDS_0387 mCherry | insert for introduction of a Sputcn32_0387-mCherry hybrid in pNTPS138-R6K | this study |
| pNPTS138-R6KT_CDS_0387 LuxCDABE | insert for introduction of a translational fusion of Sputcn32_0387-mCherry to *luxCDABE* in pNTPS138-R6K | this study |

**Supplementary Table 3: Oligonucleotides used in this study**

| Name | Construct/Purpose | Sequence |
|---|---|---|
| DE54 | FlaA1_T174C_FlaB1_T166C_S174C | GAATTCGTGGATCCAGATtgaagttaaagtgtctggga |
| DE55 | FlaA1_T174C_FlaB1_T166C_S174C | agttgcaatcgttaaACAactaaccattaaactccccg |
| DE56 | FlaA1_T174C_FlaB1_T166C_S174C | agtttaatggttagtTGTttaacgattgcaacttcagg |
| DE57 | FlaA1_T174C_FlaB1_T166C_S174C | aacttttaatgctgatgcACAggtttttgacacagaaatcgta |
| DE58 | FlaA1_T174C_FlaB1_T166C_S174C | tcagcattaaaagttggtTGTttagatattaaaggctctgctcg |
| DE59 | FlaA1_T174C_FlaB1_T166C_S174C | CAAGCTTCTCTGCAGGATctgtcacttcagataatttttcag |
| DE60 | FlaA1_FlaB1_scr | tatctagacctgaccccatgcc |
| DE62 | FlaA1_FlaB1_scr | aattttgatgcgactacccccg |
| DE01 | FlaA2_C160_ FlaB2_C156 | CAAGCTTCTCTGCAGGATGTCGCCGTCGCATTTTCG |
| DE24 | FlaA2_C160_ FlaB2_C156 | TTCCAAATTGGAGCTGGAaccGCAGAAtgtCTGGATGTGAAGTTAGGC |
| DE25 | FlaA2_C160_ FlaB2_C156 | ATCCAGACATTCTGCGgtTCCAGCTCCAATTTGGAA |
| DE26 | FlaA2_C160_ FlaB2_C156 | GCTGAAACATTGGCCGTTtgcacaACAGCTATCGATGACGCT |
| DE27 | FlaA2_C160_ FlaB2_C156 | ATCGATAGCTGTtgtgcaAACGGCCAATGTTTCAGC |
| DE10 | FlaA2_C160_ FlaB2_C156 | GCTGAAACATTGGCCGTTTGCTGTACAGCTATCGAT |
| VK184 | FlaA2_FlaB2_scr | gttaccctttggcgcatcgg |
| VK183 | FlaA2_FlaB2_scr | gtattagcttcgatcgggattgg |
| DE133 | Δ24_Δ ORF0387 | GAATTCGTGGATCCAGATccactgggcataaacctcaccc |
| DE134 | Δ24_Δ ORF0387 | ggataaagttttattatcggacagaaatgttaaaagttaaccttagtctgg |
| DE135 | Δ24_Δ ORF0387 | gataataaaactttatcccataaaaaacgccttagatggatgc |
| DE136 | Δ24_Δ ORF0387 | CAAGCTTCTCTGCAGGATagattggcaaatgcgtttatatgggcc |
| DE137 | Δ24 | ttgagatattttattatcggacagaaatgttaaaagttaaccttagtctgg |
| DE138 | Δ24 | ccgataataaaatatctcaaaacgtccaacacaggcatattga |
| DE139 | CN32_0387_scr2 | cctgccatttacgcctaaaggatagc |
| DE140 | CN32_0387_scr1 | cgccgcttagcctcgcag |
| DE198 | 0387_LuxCDABE | catatttgccctcctttataccttaaacttagccactaaagtatctaaacgatgg |
| DE199 | 0387_LuxCDABE | gctaagtttaaggtataaaggagggcaaatatgactaaaaaaatttcattcattattaacg |
| DE200 | 0387_LuxCDABE | ttttttatgggataaagtctAatcaaacgcttcggttaagctcaaagc |
| DE201 | 0387_LuxCDABE | accgaagcgtttgatTagactttatcccataaaaaacgccttagatggatg |
| DE192 | 0387Cterm-mCherry | GAATTCGTGGATCCAGATtagatgtgatccgcgctatctctgag |
| DE193 | 0387Cterm-mCherry | TTTGGAAACGCTCCCGCCtaccttaaacttagccactaaagtatctaaacgatgg |

| | | |
|---|---|---|
| DE194 | 0387Cterm-mCherry | tttaaggtaGGCGGGAGCGTTTCCAAAGGGGAAGAGGACAATATGGC |
| DE195 | 0387Cterm-mCherry | tttatgggataaagtttaTTATTTGTATAACTCATCCATACCACCAGTCGAATG |
| DE196 | 0387Cterm-mCherry | GATGAGTTATACAAATAAtaaactttatcccataaaaaacgccttagatggatg |
| DE197 | 0387Cterm-scr | cgtgctcgcgataccattaaccaattg |

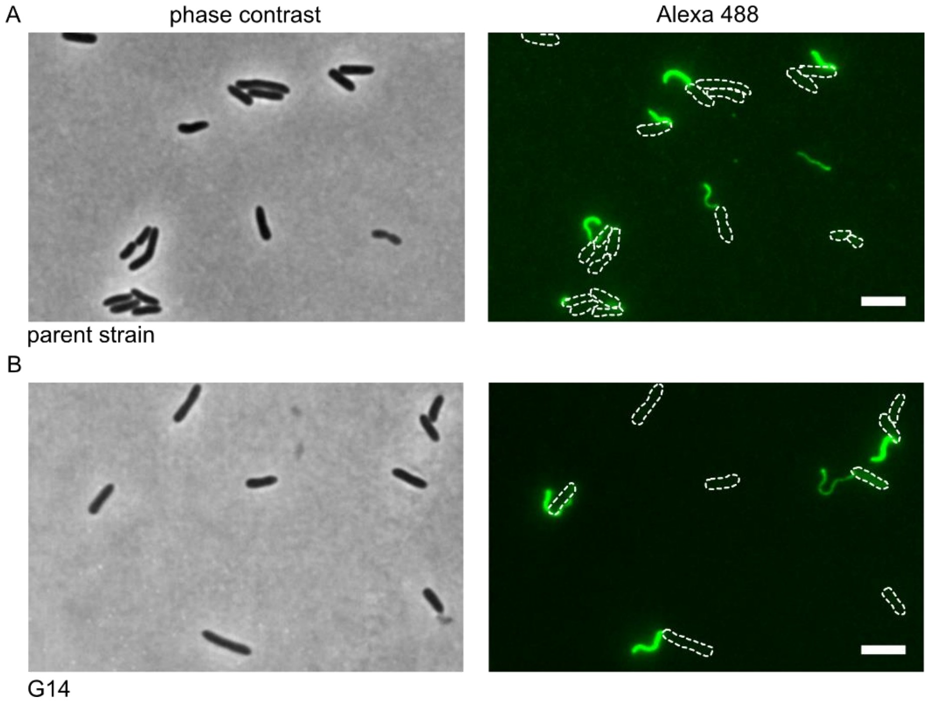

**Supplementary Figure 1: Cell morphology and flagella staining of wild-type (A) and evolved (G14; B) mutant cells.** Displayed are micrographs of cells with fluorescently labeled flagella, the cell positions are outlined in the fluorescent panels (right).

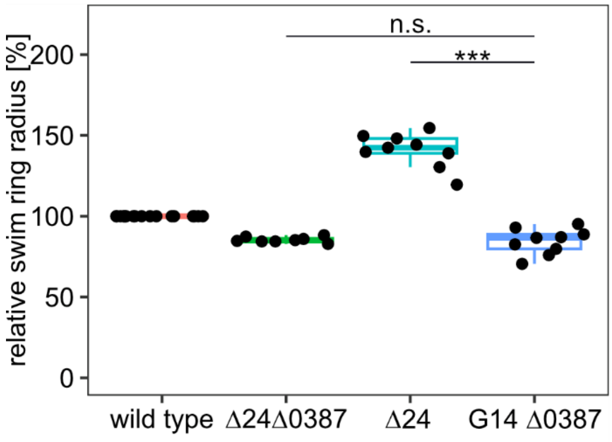

**Supplementary Figure 2: Spreading phenotype of the evolved and control strains as indicated.** The asterisks indicate the significance according to a pairwise t-test ($p < 0.01$); n.s., not significant.

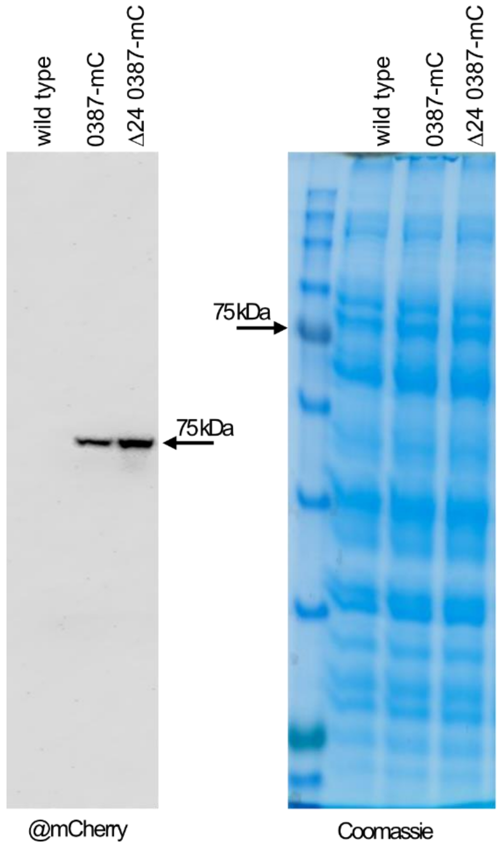

**Supplementary Figure 3: Full western and loading control (Coomassie-stained PAGE) shown in main figure 3C.** The mCherry fusion to MCP0387 results in a stable protein.

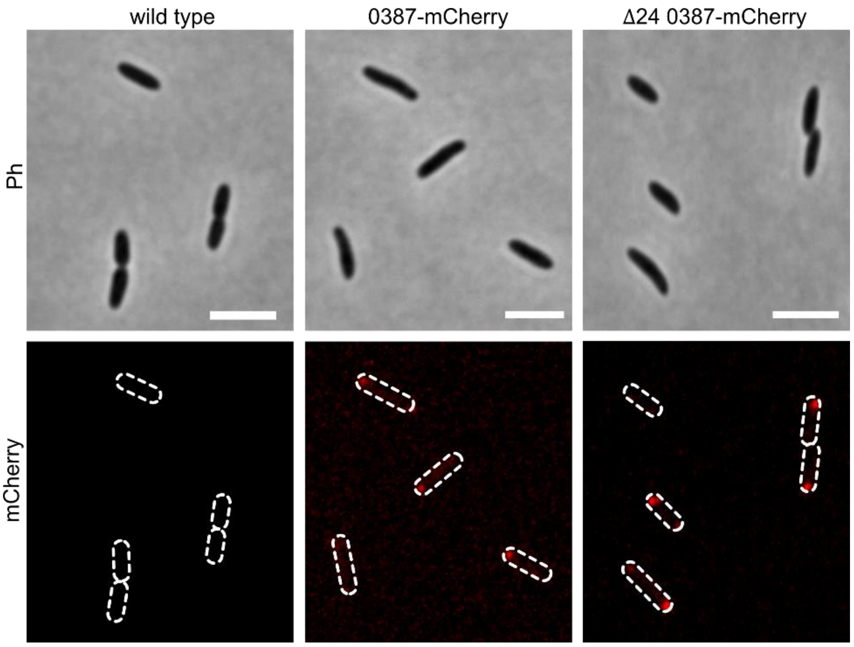

**Supplementary Figure 4: Fluorescence microscopy including the wild type shown in main figure 3D.** No signal occurs in non-tagged wild-type cells. The scale bars equal 5 µm.

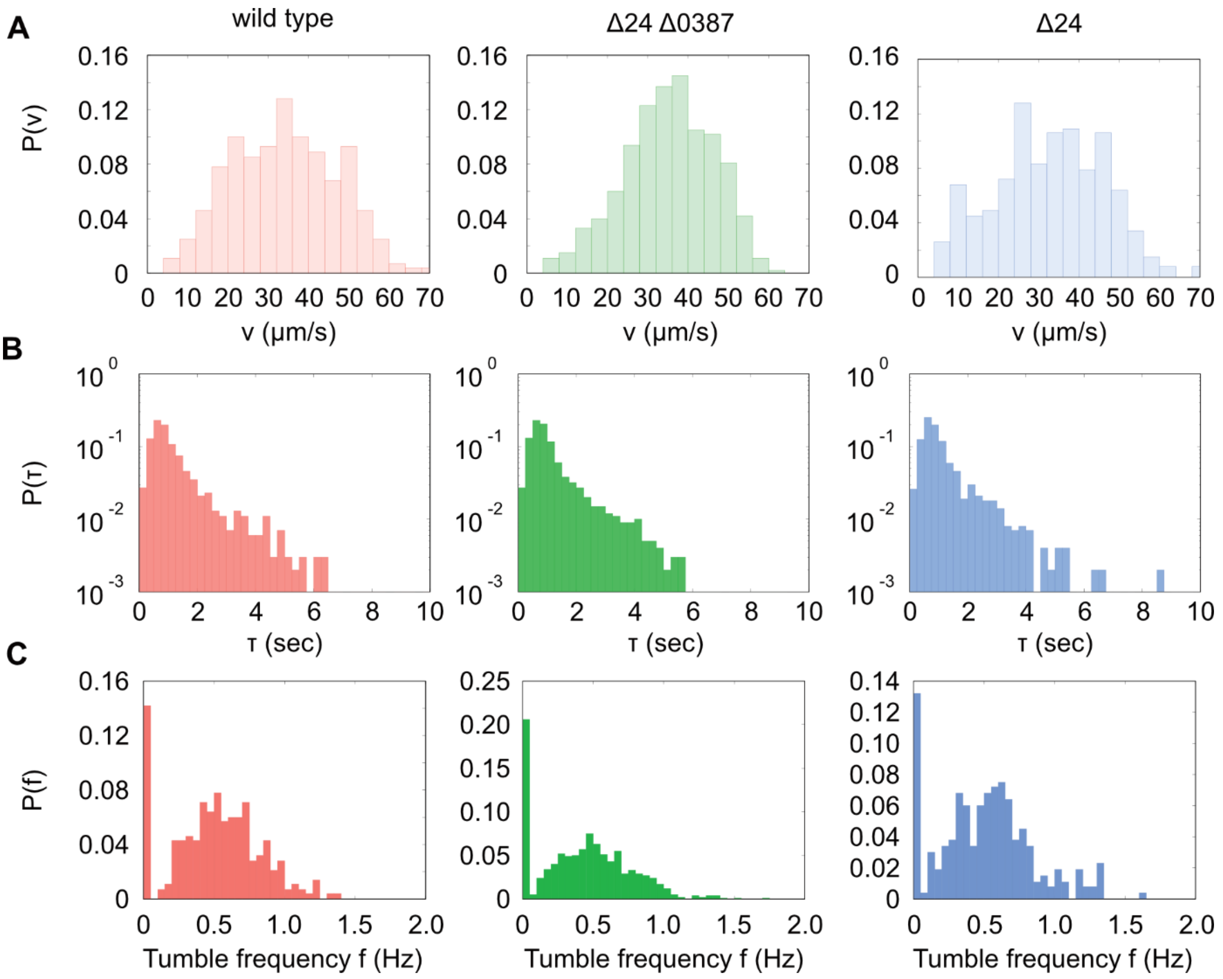

**Supplementary Figure 5: Quantification of cell tracking under planktonic conditions. A)** Distribution of cell velocities. **B)** Distribution of run lengths. **C)** Distribution of tumble frequencies. The number of tracks analyzed were 281 for the wild type, 935 for the Δ24Δ0387 mutant and 265 for the Δ24 mutant.

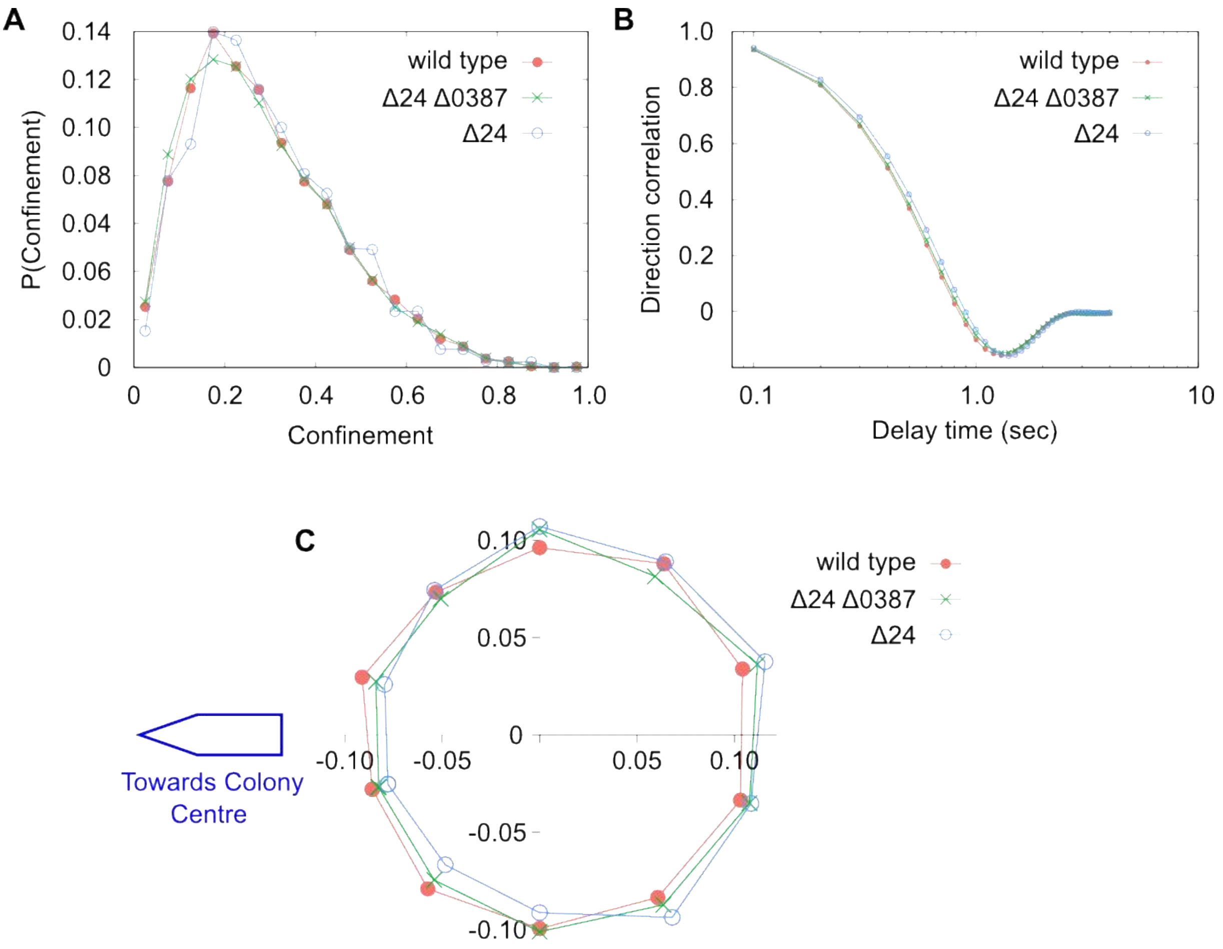

**Supplementary Figure 6: Quantification of cell tracking in soft agar. A)** Determination of the Confinement ratio, defined as $\langle t\ (\tau)\ \cdot t(0)\ \rangle$, where $t$ is the unit vector displacement between two subsequent frames. **B)** Determination of the angular distribution of steps for each strain, defined as a probability of motion in a particular direction. **C)** Each cell displacement was placed into one of ten angular bins (width 36°). All cells show a roughly isotropic distribution of displacement directions, with a slight bias away from the colony centre: the circles' centres are displaced slightly to the right of the origin. The number of tracks analyzed was 9,931 for the wild type, 11,393 for the Δ24Δ0387 mutant and 1,709 for the Δ24 mutant.

Caption Supplementary Movie 1: Time-lapse scanning recording of soft-agar spreading assays using the indicated strains. Each petri dish was inoculated with a single culture and incubated at 30°C for 18 h. The plates were scanned in 15-minutes time intervals.
